# Divergent changes in social stress-induced motivation in male and female mice

**DOI:** 10.1101/2024.10.02.616310

**Authors:** Megan McGraw, Cooper Christensen, Hailey Nelson, Ai-Jun Li, Emily Qualls-Creekmore

## Abstract

Exposure to stressors has been shown to dysregulate motivated behaviors in a bidirectional manner over time. The relationship between stress and motivation is relevant to psychological disorders, including depression, binge eating, and substance abuse; however, this relationship is not well characterized, especially in females, despite their increased risk of these disorders. Social defeat stress is a common model to study stress-induced motivation changes, however, historically this model excluded females due to lack of female-to-female aggression and unreliable male-to-female aggression. Additionally, changes in motivation are often assessed well after stress exposure ends, potentially missing or occluding changes to motivation during stress. Recently, the chronic non-discriminatory social defeat stress (CNSDS) model has demonstrated social defeat of male and female C57BL/6J mice by exposing both mice to an aggressive male CD-1 mouse simultaneously. Here we use this model to directly compare changes in the motivated behavior of male and female mice during and following chronic stress. We hypothesized that motivated behavioral responses would be dysregulated during stress and that the effects would worsen as the stress exposure continued. To monitor motivated behavior, mice had access to a Feeding Experimental Device.3 (FED3), a home cage device for operant responding. Operant responding was monitored prior to, during, and after stress by measuring nose pokes for sucrose pellets on a modified progressive ratio schedule of reinforcement. Our results demonstrated divergent behavioral outcomes between males and female mice in response to stress; where male mice increased motivated behavior during stress only, whereas female mice exhibited a decrease in motivation during and after stress. This study highlights the need to investigate the effects of stress-induced motivation over time, as well as the increased need to understand differences in the stress response in females.

**Highlights:**

- Operant behaviors were monitored continuously during and after stress exposure.
- Chronic social stress produced opposite effects on motivation in males and females.
- Susceptibility to stress only influenced outcomes on female motivated behaviors.

## 1. Introduction

Stress exerts a profound and multifaceted influence on human behavior, impacting motivation in complex and often paradoxical ways [1]. Recent studies have illuminated the intricate interplay between stress and motivational disorders, revealing that chronic stress can disrupt neural circuits critical for reward processing and decision-making [1–6]. This disruption manifests diversely in that some individuals may experience diminished motivation and anhedonia characteristic of depression, while others may exhibit heightened impulsivity and compulsive behaviors typical of substance use disorder (SUD) [7, 8], binge eating [9, 10], and anxiety disorders [11, 12]. Interestingly, rodent models of stress mirror these bidirectional effects on motivation [1, 13, 14], underscoring the translational relevance of animal research to human conditions. Despite these parallels, our understanding of the underlying mechanisms and temporal dynamics governing these disparate outcomes remains incomplete. Key questions persist regarding the role of stress duration, intensity, and individual variability in shaping these responses. Advancing our knowledge in these areas promises not only to deepen our understanding of stress-related motivational disorders but also to inform more targeted therapeutic strategies aimed at restoring adaptive motivation in affected individuals.

Rodent models of stress have begun to parse out the effects of stress on motivation. These models have included immobilization stress, uncontrollable mild stress, foot shock, and other environment-dependent stressors [1, 15, 16]. The induction of social stress has also been used to probe changes in motivation, but various models of this complex stressor have produced contradictory results across measures of motivated behaviors [7, 17, 18]. Models like chronic social defeat stress (CSDS) are used as ethological models to study the consequences of interpersonal conflict, or social stress, which is a common source of stress in humans, revealing a translational gap in our understanding of stress and motivation. Additionally, research into motivation is spread across different types of motivated behaviors. Some studies use natural rewards while others study motivation for drugs of abuse. Correspondingly, the method by which motivation is functionally measured can range from more naturalistic behaviors in rodents, like seeking behaviors, versus goal-directed methods via operant responding. Comparing studies using various combinations of these approaches reveals an intricate relationship between stress and motivation. Feeding is a constellation of behaviors necessary for survival that is sensitive to stress-induced changes and is sensitive to motivated state. Chronic stress studies have demonstrated both an overall increase in food intake [19–23] as well as an increase in high fat [6, 21, 24] and/or high sugar [6, 25] intake in response to different chronic stressors. However, other studies demonstrated either decreased standard chow, high fat and/or high sugar intake, and sucrose preference [7, 21, 22, 26–28] or no change in high fat diet intake [21, 22] during stress. However, motivation for ad libitum food intake can be altered by stress-induced metabolic changes [29], therefore models of operant behavior are important for parsing out changes in motivation from changes in free feeding behaviors. Requiring rodents to lever press or nose poke for sucrose under various reinforcement schedules has shown decreased responses for sucrose after stress [30–33], and at select time points during stress [7, 34, 35]. However, at extended time points after stress, response for sucrose was increased [18, 36]. Motivation for self-administration of substances of abuse is generally thought to be increased because of stress, simulating stress-induced relapse and craving [1, 37], however patterns are not the same across all substances and whether social stress is continuous or episodic can impact whether cocaine administration is decreased or increased [38].

As in humans, chronic stress outcomes are variable in rodents due to differences in stress paradigms, variability in aggressor activity, and individual variability. While we cannot control the exact severity of aggressor attacks, nor the individual’s perception of the stress, we can control the details of the paradigm. Over the years, there have been many variations on how to induce social stress [7, 39–46]. Without reviewing all potential variations, common differences in methods include: the duration of the model, use of a conspecific versus hyper-aggressive strains, daily time spent with an aggressor, and the level of physicality allowed before outside intervention; all of which alter the experience of the stress. Additional variability is present in the form of non-attack centric methods such as social crowding [47, 48]. These variables all contribute to the ‘inverted-U’ model of stress which states that there is a peak amount of stress that promotes arousal, and in small doses this can increase motivated behaviors, but as stress increases past a certain point, deleterious effects occur, and motivation is decreased [49]. It is important to note that these patterns are even less clear in female rodents due to difficulties inducing social defeat [50], however, researchers have begun the work to modify protocols to include females.

It is important to note that models designed to include females come with new sources of variability. Models have tried to induce male attacks on females via chemogenetic activation of discrete brain regions [51], application of male urine on the female [52], and simultaneous presentation of a male and female mouse to a male aggressor [44]. Other models have attempted to induce inter-female aggression, arguably more comparable to inter-male aggression, via the usage of maternal aggression [53] and rival aggression [42]. Alternatively, models of vicarious social defeat [54] and social crowding [47] have also been used to induce observable levels of social stress in females without the requirement of physical aggression. The models of female stress are also all subject to differences in the same variables as male-dominant protocols making it increasingly more difficult to characterize stress-motivation relationship.

In addition to the methodological variability in protocols used for social stress, there is also the aspect of timing that could account for inconsistencies in results. Nearly all studies utilizing operant behavior as an indicator of motivation for natural reward or substances of abuse are performed after the conclusion of stress [32, 33], commonly weeks after stress because the training period also occurs after stress [18, 26, 30, 36, 55–57]. This limits the time frame in which we have knowledge of stress-induced motivated behavior to well after the cessation of stress. The studies that have monitored operant responding during stress are usually limited to one middle time point [7, 31, 35] and/or monitor for time-limited sessions closely adjacent to daily stress exposure [7, 27, 31, 34, 35]. Behavioral measures in these previous studies are confined to small temporal windows that may be missing relevant behavioral changes. Studies that monitor food intake can capture daily changes in intake, which has shown that effects on food motivation are not linear during and after stress [20, 21, 29, 58]. However confounding variables such as changes in metabolic states during stress [29, 58] and accuracy of food measurement techniques [59] complicates the translation of food intake to motivation. Feeding studies are also limited as hedonic measures of motivation, thus decreasing their utility for assessing goal-directed behavior.

The limited timeframe in which we can actively monitor the motivational state of stressed rodents is due largely to technological limitations of standard behavioral testing equipment and protocols. These barriers are potentially obscuring temporal changes in motivation that could reveal important behavioral, and thus neurological, responses to chronic social stress. Additionally, measures of motivation are taken outside of the stress context, both physically and temporally, which may alter the presentation of stress responses. Operant responding commonly requires extensive training in a novel operant box separate from the home cage. This also typically occurs after the conclusion of stress, causing an extended period of time between the stress exposure and the behavioral measurement. As humans are rarely able to fully escape the stressors of daily life, as well as the contexts in which they occur, it is pivotal that we can monitor and probe behavioral changes on a broader timescale to understand the development of stress-induced disordered motivation. Recently, the Feeding Experimental Device (FED) was developed as an in-home cage device to monitor food intake continuously with minimal researcher intervention [60]. This device is equipped with nose poke ports, a variety of lights, and sound capabilities allowing it to work as a fully programmable in-home operant device. Allowing access to the FED whenever they are in the home cage eliminates the need for food restriction, which is a confounding stressor. Additionally, extended access in the home cage allows for training to be performed before stress. This creates the ability to monitor goal-directed nose poke behavior and sucrose collection on a minute-to-minute timescale throughout all phases of stress and recovery.

As a result of the limitations in temporal monitoring of motivated behaviors and the differences in social defeat protocols in males and females, it remains unclear how and when changes in stress-induced motivated behavior, whether increased or decreased, occur. In this study we sought to combine the usage of the FED3 in-home operant device and a sex-inclusive model of chronic social defeat [44], to characterize the progression of stress-induced motivated behaviors throughout the stress exposure period and into recovery from the stress. We employed a modified progressive ratio schedule [61] that allows the mice to continually earn sucrose rewards around-the-clock while still increasing the effort necessary to gain the next reward. We hypothesized that we would detect more pronounced changes in stress-induced motivated behavior during stress compared to after, and that responses would increase as stress progressed from acute to chronic.

## 2. Materials and Methods

### 2.1 Animals

All experimental procedures were approved by the Institutional Animal Care and Use Committee at Washington State University. Male and female C57BL/6J mice used for experiments were obtained from Jackson Laboratory (stock #000664). Retired male CD-1 breeders, used as aggressor mice, were obtained from Charles River (stock #022). All mice were housed at 22°C on a 12 h light/dark cycle (7AM on; 7PM off). Food (Purina lab chow 5001) and water were available ad libitum throughout all protocols. C57BL/6J mice were 12 weeks old at the beginning of stress exposure.

### 2.2 Chronic Non-Discriminatory Social Defeat Stress (CNSDS)

C57BL/6J mice remained group housed for one week following arrival for acclimation and recovery from transport. Mice were then randomly selected for control or stress conditions and pair housed with an age- and sex-matched conspecific in social subordination cages (31 L x 17 W x 15 D cm cages). Pair-housed mice were divided via a plexiglass divider perforated with holes allowing for sensory, but not physical, exposure. Cages were customized to allow continuous access to an in-home operant device (FED3) for control and stressed mice only (**Fig. 1D-E**). Stressed and control mice were housed in separate rooms to avoid exposure of control mice to stress-induced sensory cues.

**Figure 1.**
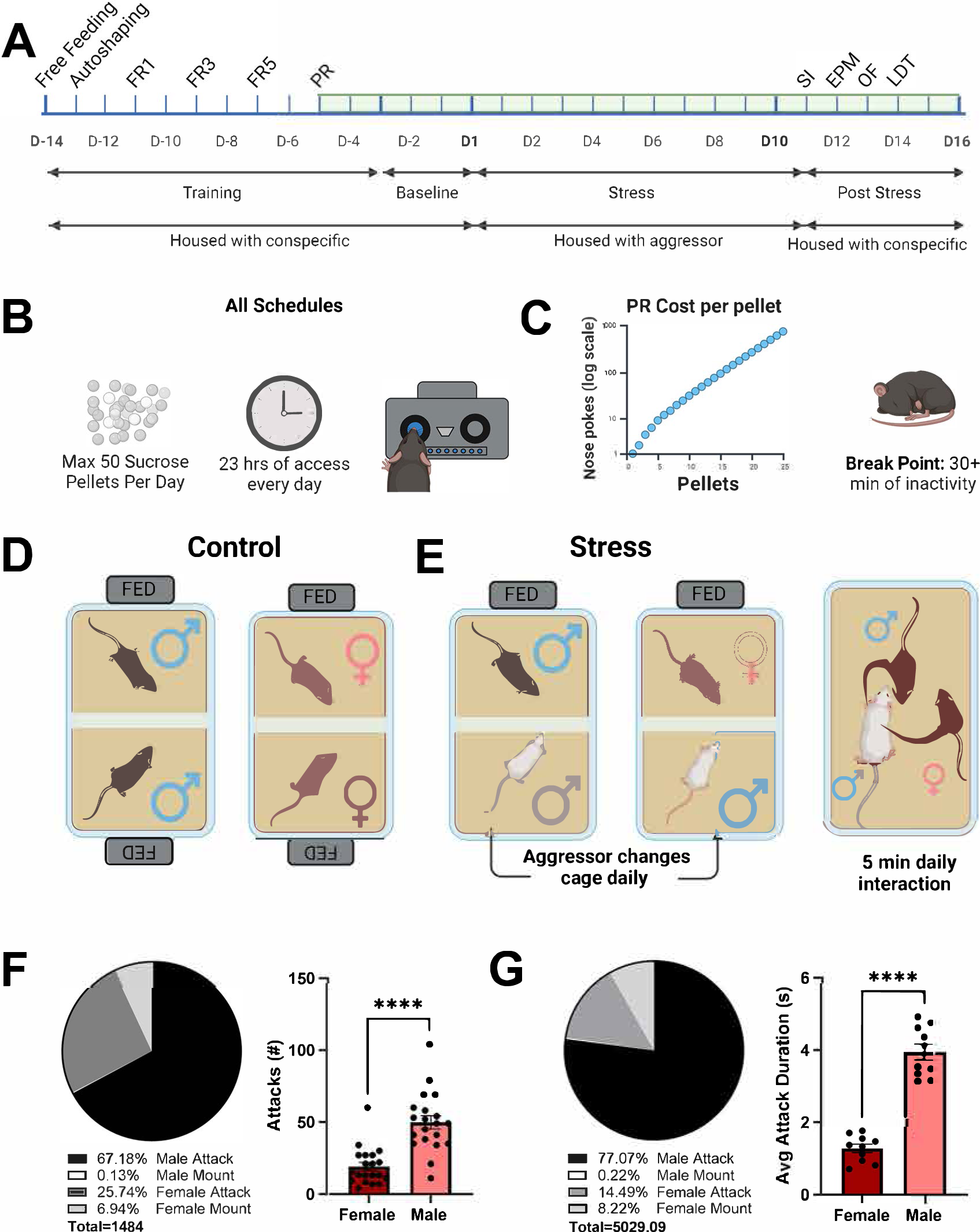
Description of experimental design and characterization of chronic non-discriminatory social defeat stress (CNSDS) model. **A.** Timeline of operant training, stress exposure, and housing conditions throughout experiment duration. **B.** Depiction of the FED restrictions during all schedules of use. On the progressive ratio schedule **(C)** the cost per pellet increased exponentially (left) until a break point was recorded due to inactivity (right). **D.** Schematic of housing conditions of control mice. **E.** Housing conditions of stressed mice during the 10 days of stress (left) and depiction of the daily 5 min physical exposure (right). **F.** Representative pie chart of the number of aggressor-initiated behaviors measured by sex (left). Males were attacked more frequently than females. **G.** Representative pie chart of the overall duration of aggressor-initiated behaviors measured by sex (left). Males were attacked for longer than females. ***p<0.001.

The chronic social stress paradigm was performed as previously described [44]. Briefly, during the 10 days of stress, male (N = 20) and female (N = 20) mice under the stress condition, hereafter referred to as experimental mice/pairs, were co-housed with retired breeder CD-1 male mice that were pre-screened for aggression towards both male and female mice [41, 44]. Mice were considered aggressive if they attacked both male and female mice separately on at least 2 out of 3 testing days within 5 minutes. Of the mice categorized as ‘aggressive’, two mice were assigned to each pair of experimental male and female mice. One CD-1 was designated to be used during the daily physical interactions, hereafter referred to as ‘aggressor’, and the other CD-1 was used for overnight sensory exposure and never physically interacted with experimental mice, hereafter referred to as ‘sensory aggressor’. On day 1, the aggressors and sensory aggressors were placed with their designated experimental mouse, replacing the non-experimental/non-control pair housed conspecific. Aggressors were initially placed in the male experimental mouse cages and acclimated for 30 minutes before the female mice were also placed in the male home cage. The divider was then removed so that the experimental pair and aggressor could physically interact for 5 minutes. Physical altercations were stopped minimally, and only when the behavior was prolonged and escalated enough to endanger the safety of the mouse. (One male was removed from the experiment on day 10 due to severe injuries on the final day.) Following the 5 minutes, the perforated divider was reinserted. The visiting experimental mouse, in this case the female, was returned to their home cage to be exposed overnight to the sensory aggressor. On day 2, the aggressor and sensory aggressor were switched and allowed to acclimate ∼30 min. The male experimental mouse was placed in the female home cage for the daily interaction and was housed overnight with the sensory aggressor. This routine was repeated for the 10-day paradigm such that the experimental mice spent 5 nights with each aggressor type. 24 hours after the final bout of stress, all aggressors were removed from all cages and replaced with the original non-experimental/non-control pair-housed conspecifics for the remaining experimental days (**Fig. 1A**). Control animals remained pair-housed with their conspecifics for the duration of the experiment and were handled daily during the same time that stressed mice received CNSDS.

### 2.4 Operant Conditioning

Operant responding on the FED (Feeding Experimental Device 3.1, Open Ephys, OEPS-7510) was used as a measure of motivated behaviors throughout and after CNSDS. The FED, a wireless pellet-dispensing device [60, 62], was equipped with a nose port on either side of the pellet dispensing magazine. The devices dispensed 20 mg sucrose pellets (5TUT, Test Diet) in response to the correct number of nose pokes in the active nose port (indicated by a blue LED light in the port). Mice were limited to a maximum of 50 pellets per day to prevent the sucrose from displacing their standard diet beyond approximately 20% of daily caloric intake and to limit potentially confounding effects of satiety and high sugar diets on outcomes of motivation. Each day, the FED was unavailable from 10AM-11AM. During this time, device maintenance, animal care, body weight collection, and stress protocols occurred, all during the light cycle for the mice. The device was then available to each mouse from 11AM until they retrieved 50 pellets, or 10AM the next day (**Fig. 1B**). In addition to sucrose rewards, mice were fed standard lab chow and water ad libitum throughout training and testing.

FED training was modified from previously published protocols [62] and outlined in Figure 1A. On day one of training (D-14) mice were trained to retrieve sucrose from the magazine. Once one pellet was removed, another pellet was dispensed (Free Feeding) until 50 sucrose pellets were retrieved. Next, (D-13) a single nose poke in either the right or left port, both illuminated blue, would trigger reward cue (lights and tone) and dispense one sucrose pellet (Auto Shaping). After that (D-11), either the left or right poke was active for each mouse, counterbalanced by sex and condition, and remained the active port for the duration of training and testing. A fixed ratio (FR) schedule was used such that a single poke in the illuminated port would trigger the reward cue and pellet (FR1). On (D-9), the ratio was increased to 3 pokes for one pellet (FR3), followed by an increase to five pokes (FR5, D-7). Finally, mice were moved to a modified progressive ratio (PR) schedule (D-5). A classic PR schedule exponentially increases the number of nose pokes necessary for each subsequent pellet until the mouse no longer continues to poke. This is usually tested for one hour; thus, the mouse is limited by time and motivation as it takes longer and longer to earn each pellet. Other studies modified the PR schedule increase in a more fashion with a ‘break point reset’ when requiring operant responding for a rodent’s entire food source [61]. For our purposes, we maintained the exponential increase in pokes per sequential pellet (pokes needed = [(5 x exp((n + 1) x 0.2)) – 5]), but we included the use of a break point reset (**Fig. 1C**). Thus, after 30 minutes of complete inactivity with the FED, the PR schedule would reset, and the next pellet would only require 1 nose poke. This ensured that all mice had the opportunity to receive the max number of pellets, but differences in motivation could still be elucidated from how much they were willing to work to earn sucrose. Mice were on the PR schedule for 2 days, prior to the beginning of baseline recording of PR behavior (D-3). Due to the familiar environment of their home cage and extended time with the FED device, mice quickly learned each task.

### 2.5 Behavioral Testing

On the days following stress, experimental and control mice were exposed to the following order of behavioral tests: Social interaction test (SI), Elevated Plus Maze (EPM), Open Field (OF), and the Light Dark Test (LDT). Each testing session began at 10AM and occurred under light conditions. All behavioral tests were recorded with IR cameras mounted directly above the testing apparatus (Balser) and analyzed with video tracking software (Noldus EthoVisionXT). Videos were checked for accuracy by a reviewer.

#### 2.5.1 Social Interaction Test

The social interaction test was used to assess the susceptibility of mice to social defeat stress [41]. Briefly, control and stressed mice explored an open field (30x30cm, Noldus PhenoTyper) for 5 minutes with a mesh cage (10 x 6 x 8 cm) in the chamber (**Fig. 6C**). For the first 2.5 minutes the mesh chamber was empty and for the second 2.5 minutes, the chamber held a novel aggressor. To determine resilience, the time spent interacting within a 2-inch zone around the mesh cage with the novel aggressor present was divided by time spent in the empty zone. SI Ratio = (Time in empty interaction zone / Time in CD-1 interaction zone). The average SI Ratio of control males and females was compared to the ratio of stressed males and females, respectively. Stressed mice with SI Ratio < Control Ratio were considered susceptible to social stress [41].

#### 2.5.2 Elevated Plus Maze

The elevated plus maze was used to assess anxiety-like behaviors resulting from stressor exposure. The EPM consists of two crossed perpendicular arms (79 x 5cm) elevated 50 cm above the ground (Stoelting). Two of the four arms are enclosed by 15 cm tall walls. At the beginning of the trial the mice were placed in the center of the maze facing the left open arm. Mice were permitted to freely explore the EPM for 5 minutes.

#### 2.5.3 Open Field Test

The open field test (OFT) was used to assess novelty-induced locomotor activity and anxiety-like approach behaviors. The OFT occurred in a 40 x 40 x 35 cm grey acrylic chamber (Stoelting). The mouse was placed in the center of the chamber and allowed to explore for 5 minutes.

#### 2.5.4 Light Dark Test

The light dark test (LDT) was employed to measure anxiety-like behaviors within the approach-avoidance conflict. The LDT chamber consisted of 40 x 40 x 35 cm plexiglass box divided in half by an opaque black wall with a 7 x 7 cm opening that allowed crossing between the two chambers. The left chamber was clear and open at the top while the right chamber was opaque black with a top to remain fully enclosed (Stoelting). The mouse was placed in the black chamber and allowed to explore for 10 minutes.

### 2.6 Behavioral Scoring

Daily interactions during CNSDS were recorded by video camera from a side view (Logitech Webcam). Attack and mounting behaviors were scored (BORIS, Friard and Gamba 2016) by at least two different individuals. Onset of attack was scored when the aggressor bit and/or lunged at the experimental mouse. Attacks ended when the aggressor diverted their attention away from the experimental mouse. Time of attacks included time the aggressor spent actively chasing an experimental male or female between biting behaviors. Scoring between raters was compared for accuracy and was averaged if total discrepancy per day between raters was less than 5 seconds, otherwise, the trial was scored a third time.

### 2.7 Corticosterone (CORT) Analysis

Tail blood was collected one week prior to stress onset and 45 min after the final stress exposure. Mice were loosely restrained in a tail vein injection platform (Fisher Scientific) for a minimal tail snip. Blood was collected via sodium heparinized capillary tubes (∼75 µL, globe scientific). Whole blood samples were left on ice for one hour and then centrifuged at 4° C, 12,000 rpm for 5 min to isolate serum. Serum was stored at −80° C until ELISA (EIA corticosterone kit, Enzo Life Sciences) analysis of corticosterone could be performed.

### 2.8 Statistical Analysis

Male and female mice were used for evaluation of motivated behaviors and behavioral assays (control = 14 mice / sex, stress = 20 mice / sex). One male mouse was excluded from all data sets due to injuries sustained on the final day of stress, no other mice were excluded from behavioral assays (control = 14 mice / sex; stress = 20 female mice and 19 male mice). For FED data, 2 female stressed mice and 3 stressed males were excluded from all FED data due to SD card corruption resulting in the loss of multiple days of data (N = 14 control mice / sex, N = 18 female mice and 16 male mice). Statistical differences were analyzed using PRISM software (GraphPad Prism 10) with analysis including paired and unpaired *t* tests, One-way ANOVA, Two-Way ANOVA, and repeated measures Two-Way ANOVA. Post hoc analysis of ANOVA used Tukey’s multiple comparisons test. A *p* value < 0.05 was considered statistically significant and notated as follows: *p < 0.05, **p < 0.01, ***p < 0.001.

## 3. Results

Previous studies have indicated the utility of CNSDS in inducing behavioral and physiological indicators of stress in both male and female mice [21, 30, 44]. To determine the effect of chronic social stress on motivated behaviors during stress in male and female mice we combined CNSDS with long term monitoring of operant behavior before, during, and after social defeat stress. Thus, we trained male (N = 34) and female (N = 34) mice to respond on a modified PR schedule for sucrose pellets and recorded baseline responding prior to stress (**Fig. 1A-C**). Then we began the CNSDS protocol where we exposed male (N = 20) and female (N = 20) experimental mice to retired CD-1 breeders for 5 minutes on 10 consecutive days (**Fig. 1A, E**). Consistent with previous studies, male mice were attacked more frequently (**Fig 1F**, unpaired *t* test, T_(32)_ = 5.647, ***p < 0.00001) and for longer durations than females (**Fig 1G**, unpaired *t* test, T_(13.79)_ = 10.78, ***p < 0.00001).

### 3.1 CNSDS induced opposing stress-related endocrine and behavioral outcomes in male and female mice

To confirm the induction of stress via CNSDS on male and female mice, we monitored daily changes in body weight (**Fig 2A-B**), corticosterone levels before and after stress (**Fig 2C, 2G**), and three separate assays of anxiety-like behavior (**Fig 2D-F, 2H-J**). Female stressed mice lost weight overall and their weight gain remained decreased compared to non-stressed controls on almost all days (**Fig. 2A**, unpaired *t* tests, N = 14 control mice, N = 18 stressed mice, *p < 0.05). Male mice overall weighed the same weight by the end of monitoring, stressed male mice temporarily gained more weight than stressed mice on days 4, 7, 8, and 9 during stress as well as 2 days following the conclusion of stress (**Fig. 2B**, unpaired *t* tests, N = 14 control mice, N = 16 stressed mice, *p < 0.05). Corticosterone levels were measured one week prior to stress (pre) and 45 - 55 min after the final day of stress (post). Post stress corticosterone levels in females were decreased compared to pre levels regardless of stress condition (**Fig 2C**, paired *t* test, N = 11 control mice, T_(10)_ = 2.71, *p = 0.02; N = 17 stressed mice, T_(16)_ = 10.04, ***p < 0.0001). While levels were similar pre stress (**Fig 2C**, unpaired *t* test, T_(26)_ = 1.84, p = 0.077), stressed female corticosterone levels were further decreased compared to control mice post stress (**Fig 2C**, unpaired *t* test, T_(26)_ = 3.07, **p = 0.005). Despite decreased levels of peripheral corticosterone, stressed females demonstrated increased anxiety like behaviors in the open field (OF) test (**Fig. 2E**, unpaired *t* test, N = 14 control mice, N = 20 stressed mice, T_(32)_ = 2.73, *p = 0.01) and the light dark box test (LDT, **Fig. 2F**, unpaired *t* test, N = 14 control mice, N = 20 stressed mice, T_(32)_ = 4.54, ***p < 0.0001). Stressed females did not differ in time spent in the open arms of the elevated plus maze (EPM) compared to control (**Fig. 2D**, unpaired *t* test, N = 14 control mice, N = 20 stressed mice, T_(32)_ = 1.467, p = 0.15).

**Figure 2.**
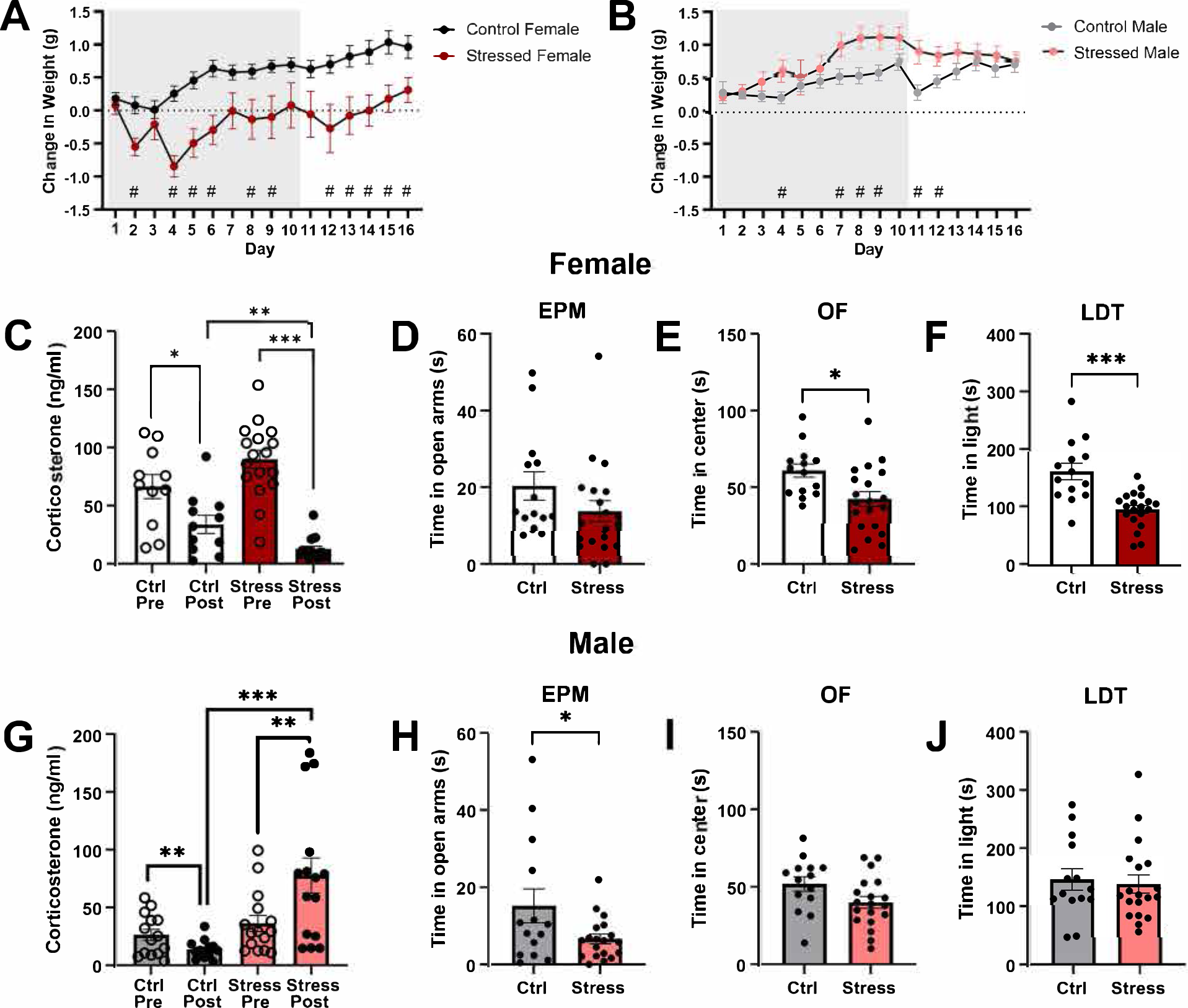
CNSDS induced divergent measures of stress in a sex specific manner. When monitoring change in body weight (g) compared to baseline, **(A)** female stressed mice lost weight compared to controls during stress, while **(B)** males gained weight during stress, but normalized after stress. **C.** Both control and stressed female corticosterone (CORT) levels were decreased compared to baseline measures. However, stressed female CORT was decreased after stress compared to controls at the same time. **D.** Stressed females spent similar amounts of time in the open arms of the Elevated Plus Maze (EPM) as controls, however they spent less time in the center of the Open Field (OF) test **(E)** and in the light chamber of the Light Dark Test (LDT) **(F)** compared to controls. **G.** Conversely, male CORT levels after stress were increased in stressed males compared to baseline levels and control animals during the same time. However, stressed males only spent less time in the open arms of the EPM **(H)** compared to controls but spent a similar amount of time in the center of the OF **(I)**, and in the light of LDT **(J)** as controls. # p<0.05, multiple unpaired t tests, *p<0.05, **p<0.01, ***p<0.001

Corticosterone levels in control males were significantly decreased at the post time point (**Fig 2G**, paired *t* test, N = 14 control mice, T_(13)_ = 3.03, **p = 0.01), however stressed mice demonstrated increased corticosterone levels after stress (**Fig. 2G**, paired *t* test, N = 15 stressed mice, T_(14)_ = 2.94, *p = 0.011). Levels of corticosterone were similar between control and stressed males pre-stress (**Fig. 2G**, unpaired *t* test, T_(27)_ = 1.16, p = 0.26), while levels were increased in stressed mice compared to control mice after stress (**Fig. 2G**, unpaired *t* test, T_(27)_ = 4.02, ***p = 0.0004). During the three behavioral assays, stressed males demonstrated increased anxiety-like behaviors in the elevated plus maze (**Fig. 2H**, unpaired *t* test, N = 14 control mice, N = 19 stressed mice, T_(31)_ = 2.13, *p = 0.04), but not the open field test (**Fig. 2I**, unpaired *t* test, N = 14 control mice, n = 19 stressed, T_(31)_ = 1.990, p = 0.0555) or light dark test (**Fig. 2J**, unpaired *t* test, N = 14 control mice, N = 19 stressed mice, T_(31)_ = 0.32, p = 0.75). While results differed by sex, both males and females demonstrated effects of stress in response to CNSDS.

### 3.2 Females have increased basal operant responding

To characterize our novel long-term operant model of motivated behaviors we first sought to monitor basal responding. We assessed the number of correct nose pokes performed to reach a max of 50 sucrose pellets on the progressive ratio schedule (**Fig. 3A**). Secondly, we monitored the number of break points occurring per day, i.e. the number of times they stopped responding for at least 30 min (**Fig. 3B**). Finally, we reported their max break point as nose pokes, i.e. the max number of nose pokes the mouse was willing to complete for a single sucrose pellet per day (**Fig. 3C**). For basal data (**Fig. 3**) both stressed and control mice were collapsed to test sex differences in basal responding before the stress protocol began. Female mice demonstrated an increase in number of nose pokes compared to males (**Fig. 3A**, unpaired *t* test, N = 32 female mice, N = 30 male mice, T_(60)_ = 3.87, ***p = 0.0003). Additionally, the number of break points occurring was decreased in females compared to males (**Fig. 3B**, unpaired *t* test, N = 32 female mice, N = 30 male mice, T_(60)_ = 4.14, ***p < 0.0001). These results are further reflected by the increased max break point in females compared to males (**Fig. 3C**, unpaired *t* test, N = 32 female mice, N = 30 male mice, T_(60)_ = 2.44, *p = 0.018). Our results reflect previous findings that female rodents have increased rates of operant responding during progressive ratio tasks for rewards compared to males [63, 64].

**Figure 3.**
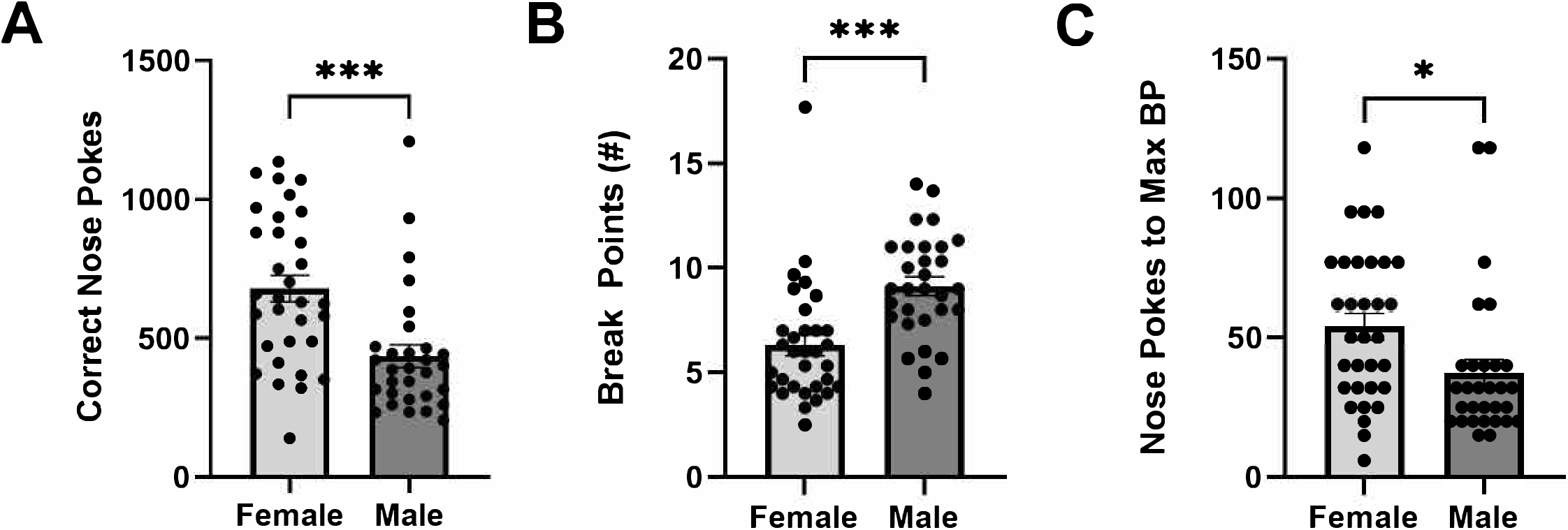
Basal levels of operant responding are increased in females. During basal conditions, female mice correctly nose poked more times per day than males for sucrose reward **(A)** and triggered less break points per day **(B)**. Additionally, the max number of pokes per pellet per day was higher in females than males **(C)**. *p<0.05, ***p<0.001

Considering increased basal responding in females, as well as individual variability in responding within sexes, all subsequent operant measures during and after stress were normalized to individual average baseline responses (**Fig. 3**). To assess overall changes in motivation during stress and recovery, daily responses were averaged into stress (Days 1-10) and post stress (Days 11-16) for each measure of operant responding.

### 3.3 Stress alters motivated behaviors in a sex-specific manner

Female stressed mice overall decreased their motivated behaviors compared to baseline activity (**Fig. 4A-C**). There was a significant effect of time (**Fig. 4A**, Two-way ANOVA, N =14 control mice, N = 18 stressed mice, F _(1.525, 45.75)_ = 24.2, ***p < 0.0001) and stress (**Fig. 4A**, Two-way ANOVA, N = 14 control mice, N = 18 stressed mice, F_(1, 30)_ = 11.16, **p = 0.002) on the average number of correct nose pokes in females. There was also a significant interaction of both time and stress (**Fig. 4A**, Two-way ANOVA, N =14 control mice, N = 18 stressed, F_(2, 60)_ = 5.98, **p = 0.004). Alternatively, there was a significant effect of time (**Fig. 4B**, Two-way ANOVA, N = 14 control mice, N = 18 stressed mice, F (1.335, 40.05) = 21.33, ***p < 0.0001), but not stress (**Fig. 4B**, Two-way ANOVA, N = 14 control mice, N = 18 stressed mice, F (1, 30) = 2.574, p = 0.12) on the average number of break points. There was no interaction between either variable (**Fig. 4B**, Two-way ANOVA, N = 14 control mice, N = 18 stressed mice, F (2, 60) = 2.40, p = 0.10). When comparing the average number of nose pokes resulting in the max break point, there was an effect of time (**Fig. 4C**, Two-way ANOVA, N =14 control mice, N = 18 stressed mice, F (1.58, 47.52) = 4.05, *p = 0.032) and stress (**Fig. 4C**, Two-way ANOVA, N =14 control mice, N = 18 stressed mice, F (1, 30) = 10.05, **p = 0.004). There was an interaction between both stress and time (**Fig. 4C**, Two-way ANOVA, N = 14 control mice, N = 18 stressed mice, F (2, 60) = 4.54, *p = 0.015). This demonstrated that the effect of stress on the female’s tendency to respond for rewards decreased during stress but continued to decline even after stress ended. While the number of breaks taken by the mice was not affected by stress, their activity per bout of responding was overall decreased.

**Figure 4.**
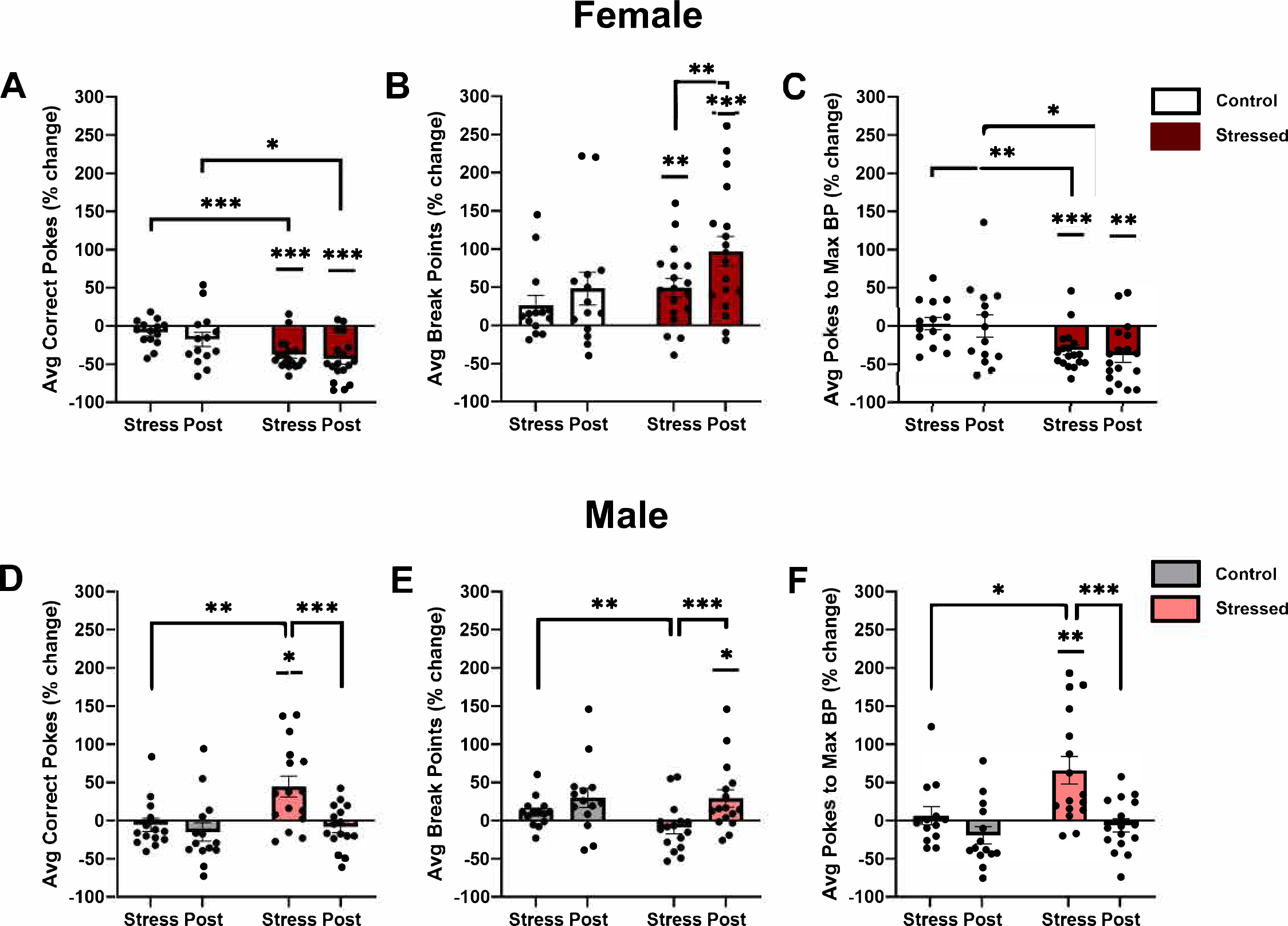
During stress, females decreased in operant responding whiles males increased. The percentage change of individual responding each day was calculated and then averaged into during and post stress groups. **A.** Female stressed mice decreased the number of nose pokes during and after stress compared to both baseline and control mice. **B.** Female stressed mice also significantly increased the number of break points over time. **C.** Consequently, they decreased their max effort for a single sucrose pellet. **D.** Conversely, stressed males increased the number of nose pokes during stress only but did increase the number of break points after stress **(E)**. Similarly, their max effort for one pellet was increased during stress, but not after **(F)**. *p<0.05, **p<0.01, ***p<0.001

In males, responding during stress was increased compared to control animals (**Fig. 4D-F**). When evaluating the average number of correct pokes in males, we found an effect of time (**Fig. 4D**, Two-way ANOVA, N =14 control mice, N = 16 stressed mice, F (1.875, 52.49) = 8.01, **p = 0.0012) and stress (**Fig. 4D**, Two-way ANOVA, N = 14 control mice, N = 16 stressed mice, F (1, 28) = 5.003, *p = 0.034). There was an interaction between stress and time (**Fig. 4D**, Two-way ANOVA, N = 14 control mice, N = 16 stressed mice, F (2, 56) = 6.020, **p = 0.004). There was an effect of time (**Fig. 4E**, Two-way ANOVA, N =14 control mice, N = 16 stressed mice, F (1.453, 40.67) = 12.38, ***p = 0.0003), but not stress (**Fig. 4E**, Two-way ANOVA, N =14 control mice, N = 16 stressed mice, F (1, 28) = 0.75, p = 0.39), on the average number of break points in males. Additionally, there was no interaction of time or stress (**Fig. 4E**, Two-way ANOVA, N = 14 control mice, N = 16 stressed mice, F (2, 56) = 1.49, p = 0.24). When comparing the average number of pokes to the max break point, there was an effect of time (**Fig. 4F**, Two-way ANOVA, N = 14 control mice, N = 16 stressed mice, F (1.622, 45.41) = 13.62, ***p < 0.0001) and stress (**Fig. 4F**, Two-way ANOVA, N = 14 control mice, N = 16 stressed mice, F (1, 28) = 5.693, *p = 0.024). There was an interaction between both variables (**Fig. 4F**, Two-way ANOVA, N = 14 control mice, N = 16 stressed mice, F (2, 56) = 5.000, *p = 0.01). Similarly to females, this demonstrated that the tendency to respond on the FEDs was not altered by stress, but that levels of activity during individual bouts was changed. During stress this effort is increased in males, while females had the opposite response and decreased effortful behavior.

The above results sought to analyze the overall changes in motivated behavior during stress and recovery, however, there remained interest in the daily changes in motivation that may become exacerbated as stress continued to occur. To assess this, we performed repeated measure assessments on motivated behavior during the stress period only (**Fig. 5**). When looking at the daily change in correct nose pokes, females had a main effect of time (**Fig. 5A**, Mixed-effects model, N = 14 control mice, N = 18 stressed mice, F (3.671, 104.8) = 2.538, *p = 0.049), stress, (**Fig. 5A**, Mixed-effects model, N = 14 control mice, N = 18 stressed mice, F (1, 30) = 18.38, ***p = 0.0002), and an interaction between both (**Fig. 5A**, Mixed-effects model, N = 14 control mice, N = 18 stressed mice, F (9, 257) = 4.26, ***p < 0.0001). Post hoc analysis demonstrated that stressed females had decreased nose pokes compared to control animals on days 2-4 and 7-10 (**Fig. 5A**, Tukey’s multiple comparisons test, *p < 0.05). Additionally, while there was no change over time in control animals nose pokes, stressed animals nose pokes moderately normalized on day 6 which was significantly increased compared to percent change in pokes on days 7 and 9 (**Fig. 5A**, Tukey’s multiple comparisons test, *p < 0.05). In males, there was no main effect of time (**Fig. 5B**, Mixed-effects model, N = 14 control mice, N = 16 stressed mice, F (2.243, 61.31) = 2.69, p = 0.07), but there was a main effect of stress (**Fig. 5B**, Mixed-effects model, N = 14 control mice, N = 16 stressed mice, F (1, 28) = 8.906, **p = 0.006). There was no interaction between the two factors (**Fig. 5B**, Mixed-effects model, N = 14 control mice, N = 16 stressed mice, F (9, 246) = 1.616, p = 0.11). Post hoc analysis showed that the main effect of stress was driven by the initial 3 days of stress, as well as days 6, 8, and 10 (**Fig. 5B**, Tukey’s multiple comparisons test, *p < 0.05).

**Figure 5.**
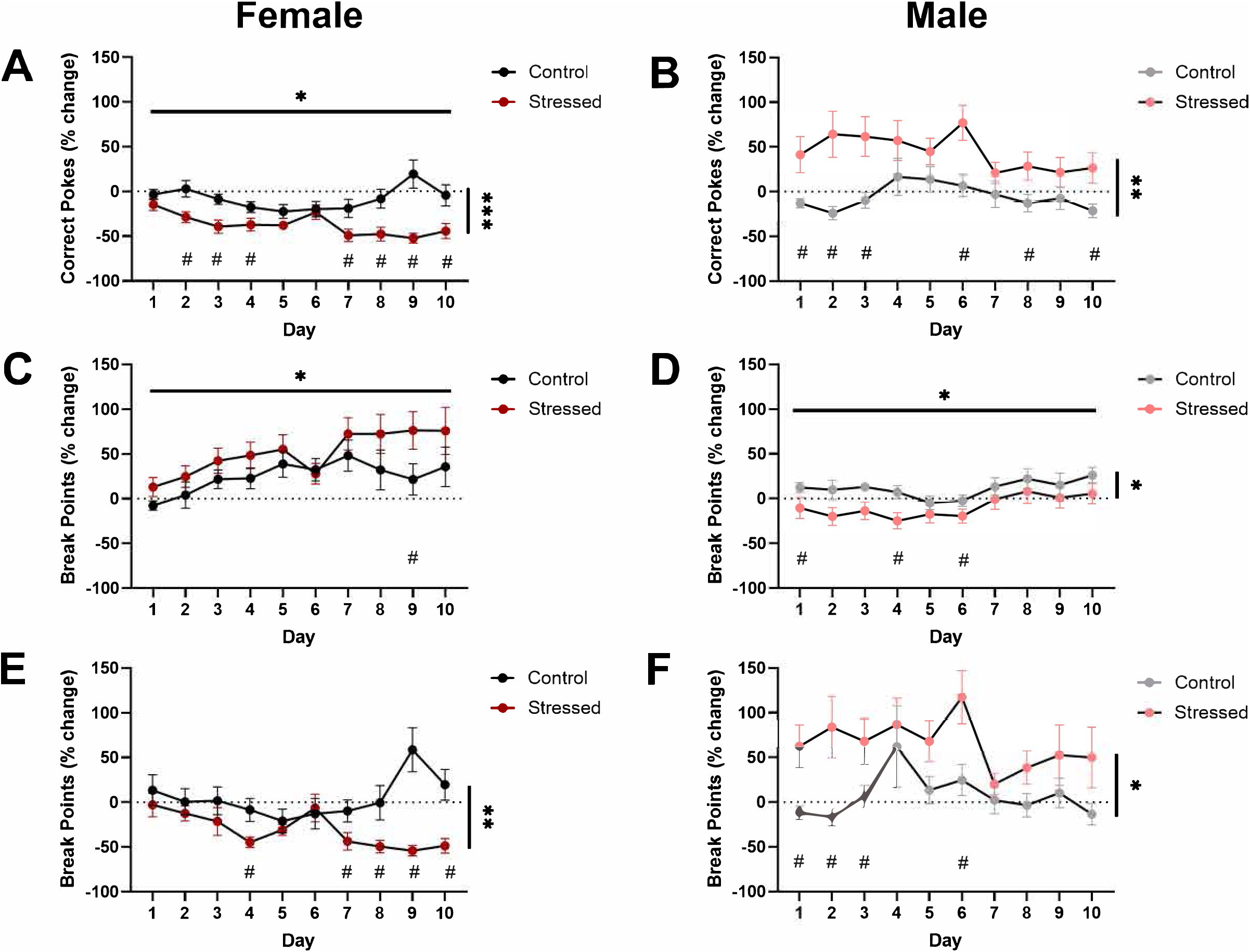
Stress-induced motivated behavior changes were not compounded by the chronic nature of the stress protocol. **A.** Female stressed mice decreased their number of pokes compared to controls during stress, but both groups changed responding over time. **C.** There was, however, no effect of stress on the number of break points over the 10 days of stress. **E.** Female max effort was decreased in stressed mice but did not worsen over time. Male however had an effect of stress in number of nose pokes **(B)**, breakpoints **(D)**, and max effort for one pellet **(F)**. # p<0.05, tukey’s multiple comparisons test, *p<0.05, **p<0.01, ***p<0.001

**Figure 6.**
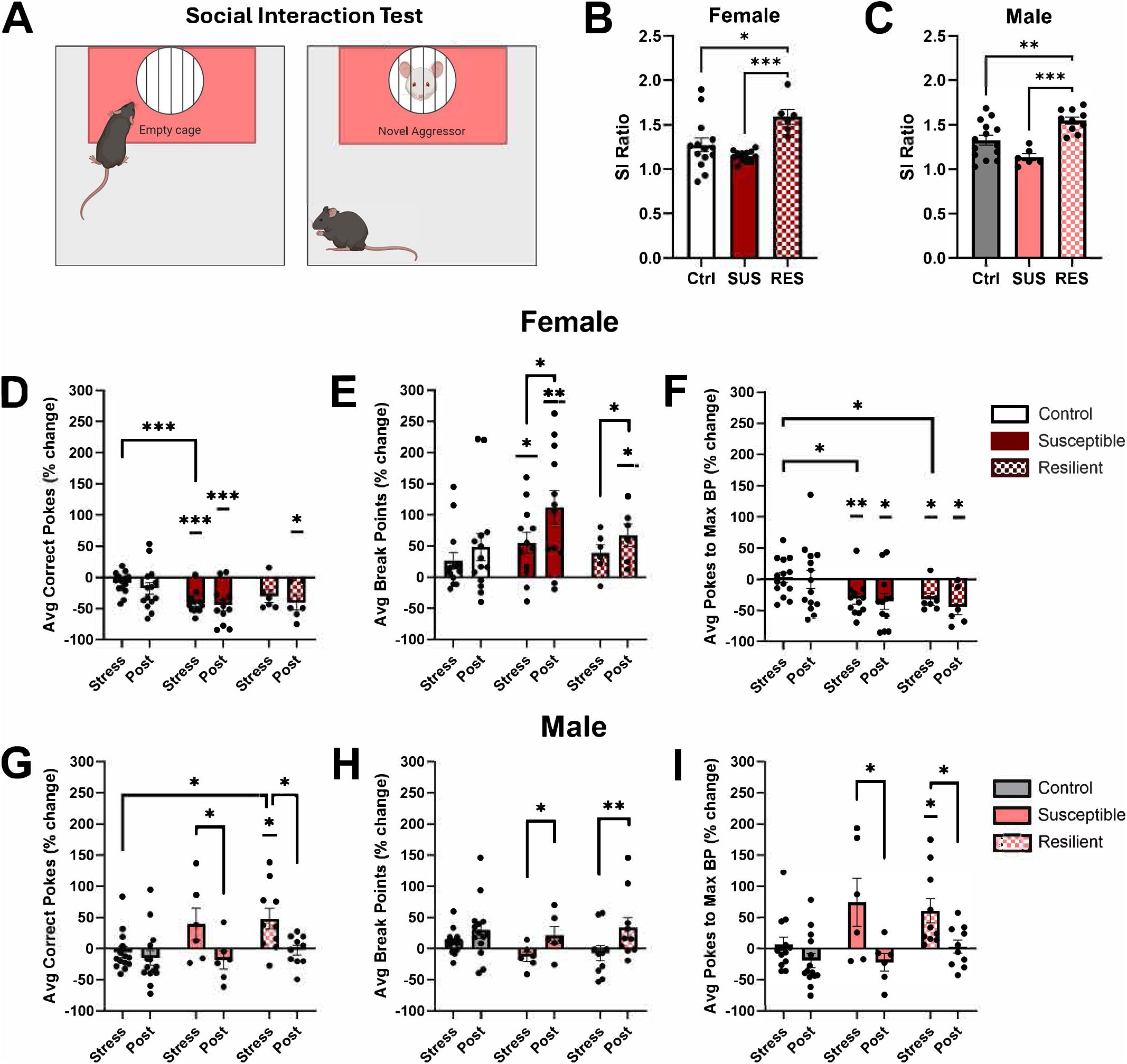
Susceptibility to social defeat stress did not robustly influence changes in stress-induced motivated behaviors. **A.** Depiction of the social interaction test. Both female **(B)** and male **(C)** mice deemed resilient (RES) to social avoidance had a higher social interaction ratio than control and susceptible (SUS) mice. Female RES and SUS mice did not differ significantly in percent change of average nose pokes **(D**), break points **(E)**, or max effort for one pellet **(F).** Similarly, male RES and SUS mice did not differ significantly in percent change of average nose pokes **(G)**, break points **(H)**, or max effort for one pellet **(I)**. *p<0.05, **p<0.01, ***p<0.001

Evaluation of the change in number of break points per day of stress in females showed a main effect of time (**Fig. 5C**, Mixed-effects model, N = 14 control mice, N = 18 stressed mice, F (2.348, 67.04) = 4.28, *p = 0.013), but not stress (**Fig. 5C**, Mixed-effects model, N = 14 control mice, N = 18 stressed mice, F (1, 30) = 1.822, p = 0.19), nor an interaction of stress and time (**Fig. 5C**, Mixed-effects model, N = 14 control mice, N = 18 stressed mice, F (9, 257) = 1.29, p = 0.24). Further demonstrating that the frequency of FED usage in females was not changed by stress. Post hoc analysis shows that in both control and stressed animals, break points are increased at day 7 showing a potential effect of familiarity and learning rather than an effect of stress. In males, however, there was an effect of time (**Fig. 5D**, Mixed-effects model, N = 14 control mice, N = 16 stressed mice, F (2.669, 72.97) = 3.77, *p = 0.018), and stress (**Fig. 5D**, Mixed-effects model, N = 14 control mice, N = 16 stressed mice, F (1, 28) = 4.273, *p = 0.048); however there was no interaction effect (**Fig. 5D**, Mixed-effects model, N = 14 control mice, N = 16 stressed mice, F (9, 246) = 0.457, p = 0.90). Thus, control and stressed males follow similar patterns of change over time, but stressed males had less break points on days 1, 4, and 6 (**Fig. 5D**, Tukey’s multiple comparisons test, *p < 0.05).

Finally, the percent change on nose pokes to the daily max break point in females was not altered by time (**Fig. 5E**, Mixed-effects model, N = 14 control mice, N = 18 stressed mice, F (4.44, 127.2) = 2.09, p = 0.08), but there was a main effect of stress (**Fig. 5E**, Mixed-effects model, N = 14 control mice, N = 18 stressed mice, F (1, 30) = 12.05, **p = 0.002) and an interaction between time and stress (**Fig. 5E**, Mixed-effects model, N = 14 control mice, N = 18 stressed mice, F (9, 258) = 4.539, ***p < 0.0001). Despite changes in nose pokes as early as day 2 (**Fig. 5A**), changes to break point values were not decreased until day 4, and then from day 7-10 (**Fig. 5E**, Tukey’s multiple comparisons test, *p < 0.05). However, these effects were not further decreased as a function of time. In males, there was no main effect of time (**Fig. 5F**, Mixed-effects model, N = 14 control mice, N = 16 stressed mice, F (3.073, 83.66) = 2.50, p = 0.064), but there was an effect of stress (**Fig. 5F**, Mixed-effects model, N = 14 control mice, N = 16 stressed mice, F (1, 28) = 7.03, *p = 0.013). There was no interaction between stress and time (**Fig. 5F**, Mixed-effects model, N = 14 control mice, N = 16 stressed mice, F (9, 245) = 0.88, p = 0.54).

Overall, these results suggested that the changes in motivation between day 1 and day 10 were not significant. Thus, the chronic effect of CNSDS did not appear to compound the severity of motivated behavioral changes each day. Rather, the experience of the stressor each day maintained either decreased (females) or increased (males) responding compared to non-stressed controls throughout the paradigm.

### 3.4 Resilience to stress does not alter patterns of stress-induced motivated behavior

Susceptibility and resilience to stress has become an important factor in understanding differences in individual responses to stress [41, 65]. CNSDS, like many social defeat paradigms, has been shown to induce susceptible and resilient populations in males and females [2, 44]. We too saw this phenomenon in our mice using the social interaction test (**Fig. 6A**. In females, our control group had an average SI ratio = 1.27. Of our stressed mice, 12 had a ratio < 1.27 and were thus considered susceptible (SUS), while the other 6 had an SI ratio > 1.27 and were considered resilient (RES) for our purposes. There was an effect of condition on the SI ratio (**Fig. 6B**, One-way ANOVA, N = 14 control mice, N = 12 SUS mice, N = 6 RES mice, F (2, 29) = 8.617, **p = 0.0012), with the RES group having a significant increase SI ratio compared to control (**Fig. 6B**, Tukey’s multiple comparisons test, T (29) = 4.299, *p = 0.013) and RES (**Fig. 6B**, T (29) = 5.86, ***p = 0.0008). In males, the average SI ratio of controls was 1.32. Of the stressed males, 6 mice were considered SUS and 10 were considered RES. There was an effect of condition on SI ratio (**Fig. 6C**, One-way ANOVA, N = 14 control mice, N = 6 SUS mice, N = 10 RES mice, F (2, 27) = 12.08, ***p = 0.0002). Again, RES mice had an increased SI ratio compared to controls **(Fig. 6C**, Tukey’s multiple comparisons test, T (27) = 4.530, **p = 0.009) and SUS mice (**Fig. 6C**, T (27) = 6.779, ***p = 0.0002).

The measures of motivated behaviors were broken down into control, SUS, and RES, based on SI ratio. This allowed us to assess whether resilience or susceptibility altered stress-induced motivated behaviors. In females, the percent change in nose pokes was still altered by time (**Fig. 6D**, Two-way ANOVA, N =14 control mice, N = 18 stressed mice, F (1.500, 43.50) = 26.53, ***p < 0.0001), condition (**Fig. 6D**, Two-way ANOVA, N = 14 control mice, N = 18 stressed mice, F (2, 29) = 5.68, **p = 0.008) and an interaction between the two (**Fig. 6D**, Two-way ANOVA, N = 14 control mice, N = 18 stressed mice, F (4, 58) = 3.052, *p = 0.024). Post hoc analysis showed that the condition effect was driven by a decrease in nose pokes during stress in the SUS mice compared to controls (**Fig. 6D**, Tukey’s multiple comparisons, T (22.9) = 6.39, **p = 0.003). However, there was no change between control and RES nor SUS and RES, demonstrating no significant difference between the RES versus SUS condition. There continued to be only a time effect on the number of break points (**Fig. 6E**, Two-way ANOVA, N =14 control mice, N = 18 stressed mice, F (1.347, 39.05) = 20.69, ***p < 0.0001), but not condition (**Fig. 6E**, Two-way ANOVA, N = 14 control mice, N = 18 stressed mice, F (2, 29) = 1.75, p = 0.19) or interaction of the two (**Fig. 6E**, Two-way ANOVA, N = 14 control mice, N = 18 stressed mice, F (4, 58) = 1.71, p = 0.16). Finally, for the nose pokes to max break point, there was a main effect of time (**Fig. 6F**, Two-way ANOVA, N = 14 control mice, N = 18 stressed mice, F (1.587, 46.01) = 6.74, **p = 0.005) and condition (**Fig. 6F**, Two-way ANOVA, N = 14 control mice, N = 18 stressed mice, F (2, 29) = 4.911, *p = 0.015), but not an interaction (**Fig. 6F**, Two-way ANOVA, N = 14 control mice, N = 18 stressed mice, F (4, 58) = 2.263, p = 0.07). Post hoc analyses show that both SUS and RES mice had similar patterns where both time points of each condition were decreased compared to baseline and were both decreased compared to control animals during stress (**Fig. 6F**, Tukey’s multiple comparisons test, *p < 0.05). However, there were no differences between SUS and RES further solidifying that in females there was no robust effect of resilience on motivated behaviors.

In males, there was an effect of time (**Fig. 6G**, Two-way ANOVA, N = 14 control mice, N = 16 stressed mice, F (1.864, 50.34) = 11.23, ***p = 0.0001), but not condition (**Fig. 6G**, Two-way ANOVA, N = 14 control mice, N = 16 stressed mice, F (2, 27) = 2.678, p = 0.09) on the number of nose pokes. There was an interaction between the two (**Fig. 6G**, Two-way ANOVA, N = 14 control mice, N = 16 stressed mice, F (4, 54) = 3.051, *p = 0.024). However, both SUS and RES mice had decreased average of nose pokes post stress compared to stress (**Fig. 6G**, Tukey’s multiple comparisons, *p < 0.05), demonstrating similar patterns between SUS and RES that are different from control. Continuing the pattern, there was only a main effect of time (**Fig. 6H**, Two-way ANOVA, N = 14 control mice, N = 16 stressed mice, F (1.456, 39.31) = 11.41, ***p = 0.0005) on number of break points, but not condition (**Fig. 6H**, Two-way ANOVA, N = 14 control mice, N = 16 stressed mice, F (2, 27) = 0.494, p = 0.62) or an interaction (**Fig. 6H**, Two-way ANOVA, N = 14 control mice, N = 16 stressed mice, F (4, 54) = 0.8188, p = 0.52). Finally, there was an effect of time on nose pokes to max break point (**Fig. 6I**, Two-way ANOVA, N = 14 control mice, N = 16 stressed mice, F (1.602, 43.25) = 18.72, ***p < 0.0001), but not condition (**Fig. 6I**, Two-way ANOVA, N = 14 control mice, N = 16 stressed mice, F (2, 27) = 2.788, p = 0.08). There was an interaction between time and condition (**Fig. 6I**, Two-way ANOVA, N = 14 control mice, N = 16 stressed mice, F (4, 54) = 3.018, *p = 0.026), but, again, this was driven by similar patterns in SUS and RES that differed from control animals. Overall, motivated behaviors monitored daily were not sensitive to differences in the susceptibility or resilience to stress. Instead, these behaviors were changed by the exposure of stress rather than factors that contribute to generalized aversion to aggressive CD-1 mice in the social interaction test.

## 4. Discussion

Our results demonstrate that goal-directed behavior can be monitored throughout a chronic social defeat paradigm, and that there are effects of time and sex in stress-induced motivated behavioral changes. While we did not find evidence to support our hypothesis that motivation would be further dysregulated over the 10 days of stress, we did find that changes in motivation during stress were significantly different from motivational changes after stress. This emphasizes the importance of investigating changes that occur during stress versus outside of stress. Additionally, we demonstrated both basal and stress-induced sex differences in motivated behaviors. Our baseline measurements confirmed that, under non-stressed conditions, female mice exert more effort on a modified progressive ratio schedule for sucrose pellets than males. When stressed, however, females significantly decrease their PR responding while males increased responding. These results overall highlight the importance and feasibility of temporally expanding motivated behavioral monitoring in both male and females simultaneously and suggest fundamental differences in behavioral coping between sexes. Here we discuss how our results fit within, and add to, our current understanding of social stress-induced motivated behavioral changes within the context of timing and sex.

### 4.1 Temporal changes of motivation associated with stress

We hypothesized that the changes in motivation would be most pronounced during days of stress, thus we utilized the FED in-home operant chamber that allowed us to do all training and testing in the home cage prior to stress. Monitoring operant responding before stress allowed for establishing individual baseline behavior, and following changes to behavior during stress, and for a week after stress while we also performed typical behavioral assays. These time frames accounted for previously under investigated windows that may hold critical behavioral changes that may point to neurophysiological underpinnings of disorders of motivation.

While we are not the first to capture changes in motivation during stress, this is the only study to our knowledge that monitors motivation during every day of stress in a non-invasive, around-the-clock manner. In doing this, we observed a decrease in female motivation during stress and an increase in motivated behavior in males based on increased nose pokes and the max effort performed for a single sucrose pellet. The finding in males is contradictory to the handful of studies that have included one or more measurements of motivation during stress timepoints. (There were no studies in females that monitored motivated behaviors during stress.) In these studies, they found either decreased motivation [7, 27, 34, 35], or no change during stress [31, 35, 36] in males. These protocols are limited to 1 hour or less of testing operant responding usually, the morning before the next stress exposure. By only monitoring for one hour and in some cases requiring responses within discreet timed trials following a cue to measure motivation [34, 35], changes in motivation may be affected by circadian influences. Chronic social defeat stress, as well as most other stressors, disrupt both quality and duration of sleep [66, 67]. Disruption of vigilance and arousal may disproportionally hinder response time in stressed mice. Additionally, energy state [68] and basal motivated behaviors [69] follows a circadian pattern and measuring operant responding in only the light cycle (in-active phase) or dark cycle (active phase) may result in divergent effects on motivated behaviors. While we cannot rule out that energy states play a role in our results, we observed that the daily number of times they interacted with the FEDs did not change as an effect of stress. However, the activity during these bouts of activity is altered with a decreased or increased response rate per bout in females and males, respectively. Additionally, our study captures motivation states immediately following stress up until they reach a max of 50 pellets each day. This means that much of our data is collected prior to their next light cycle when other studies have typically measured motivated behaviors. Thus, we likely don’t capture changes in motivation that may be a result of anticipating the next stress event, which could also explain differences in our results compared to others. These findings highlight the potential oscillating nature of stress throughout the day and the need to capture a more wholistic view of behavior in response to stress. By doing so, we confirmed that stress-induced motivation changes can occur as early as hours after the initial stress and do not appear to be worsened over 10 days of stress exposure.

Multiple studies have reported decreased goal-directed behaviors following chronic social stress in males [27, 30–33, 35, 57, 70] and females [26, 30]. With a few exceptions [32, 33] [31, 35], training for motivated behavior protocols occurs after stress. The use of the FED device allowed us to complete the operant training schedule prior to stress exposure. Thus, our work has examined stress-induced motivation in a separate time window than has been typically performed. In comparison to previous work, our findings that motivation was decreased after stress in females is consistent, despite the difference in timing [26, 30]. However, our results did not support a decrease in male motivated behaviors, but rather an increase which was specific to the stress period. Given that we measured motivated behavior during and directly following stress, our data begin to fill in the missing window of most other studies. Our results in males demonstrated an initial increase in motivation during stress that was then attenuated after stress. In the few studies which completed training for motivated behavior tasks prior to stress and were thus able to test these behaviors during or soon after stress exposure [31–33, 35], several key differences stand out. Each of these studies employed a 15-day stress protocol using the male-to-male model of social defeat stress; therefore, the duration of stress is longer, and the interaction may be more intense given the one-to-one social dynamic compared to our one-to-two CNSDS model. Another key difference in these studies is the use of food restriction during training for behavioral tasks. Food restriction is a stressor that may influence homeostatic and motivational drive. Further, one study [31] uses saccharin in their measure of motivated behavior, indicating that the caloric value of the reward may be a factor in the outcome of stress on motivation. Indeed, studies have shown that intake of calorically dense foods or rewards can buffer the stress response [25, 71, 72]. Finally, in comparison to our study, each of these studies evaluates motivated behavior using different measures of motivation that are employed at different days and time windows relevant to stress exposure. Future studies using our protocol could be used to determine how these types of methodological differences alter the development of motivated disorders on an extended timescale.

### 4.2 Complexities of sex differences in motivated behavioral changes

Considering the limited studies that included females in measurements of motivated behavior following stress reported similar effects in males and females [26, 30], we did not anticipate a sex-specific difference. However, the divergent effect we saw in males and females during and after stress exposure does support the increased incidence of stress-induced affective disorders in women [73–78]. A significant immediate and sustained decrease in motivated behavior in response to stress aligns with a stress induced depressive-like behavior that is a proxy for Major Depressive Disorder. Additionally, the increase in motivated behavior of male mice aligns with the historic increased incidence of substance use disorder in males [79, 80]. However, as attention to sampling bias and environmental factors around gender change, this margin of difference between the sexes has begun to narrow [81–83]. While these are exciting results and demonstrate models by which to continue studying these effects in males and females, these divergent results must be considered carefully with respect to the model itself.

In this model, the male and female pairs are exposed to the same physical aggressor and sensory aggressor for the same duration of time. This model is unique in that aspect, and this does add a factor of comparability between males and females that is lacking when either A) the male and female protocols are designed differently e.g. using social crowding [47], maternal aggression [84], or rival aggression [42] or B) an additional manipulation is required to induce aggression toward females but not males, such as the application of male urine on the female [52] or chemogenetic activation of aggression centers of the brain [51]. Despite significant parallel components in the CNSDS model between males and females, these mice do not experience the same stressor, irrespective of individual variability in stress perception. In both our hands, and the original model, the male mice are attacked more frequently and for a longer duration than the females. This adds a unique element of females witnessing physical aggression in addition to experiencing it. Studies have demonstrated that vicarious social defeat, or witness defeat, is a significant stressor [54, 55, 85–87]. There have been studies to compare the behavioral responses to witness defeat and chronic social defeat stress in male and female mice and rats [88]. Briefly, witness defeat more consistently alters behaviors related to social interaction scores, despair behaviors, and decreased pleasure (anhedonia). Specifically in studies using witness defeat to model social defeat in females, they have found increased measures of despair and anhedonia as measured by the sucrose preference test on the last four days of stress exposure [85]. These results complement our findings of decreased motivation for a sucrose reward during CNSDS stress. While there are limited studies of operant behavior following vicarious social defeat, especially in females, one study found an increase in ethanol intake in stressed mice, but no change in break point on a PR schedule [55]. Additionally, when comparing witness defeat versus physical defeat in males, only those who experienced witness defeat decreased their preference for sucrose 4 days after chronic stress exposure [89]. This suggests that vicarious social defeat may have variable effects on motivated behavior compared to the stress of direct aggression. Thus, the herein described effects of CNSDS may be inclusive of aspects of vicarious social defeat and direct social defeat, resulting in a more complex, and potentially more intense, stressor in females than males.

In addition to divergent changes in motivated behaviors in males and females, we also observed divergent effects of corticosterone level in response to stress. In line with previous findings, we observed an increase in corticosterone levels in males following chronic social stress [43, 44, 90, 91]. However, our robust decrease in female corticosterone levels is contrary to the results of the original CNSDS paradigm paper [44]. While increased corticosterone is traditionally seen as the hallmark of the stress response, a robust decrease of hypothalamic-pituitary-axis activity is also an effect of stress, commonly seen in individuals with post-traumatic stress disorder [92]. Indeed, the behavioral assays and change in motivated behavior suggest that our females are indeed stressed. While less common, a decrease in corticosterone following repeated stress is not unheard of. A decrease in female corticosterone levels was observed 30 minutes after the fourth episode of intermittent vicarious social defeat stress [86]. In another study, following 21 days of restraint stress, female mice had decreased levels of corticosterone compared to both baseline and 24 hours after the initial restraint [93]. These findings were accompanied by a decrease in sucrose preference, which may be relevant to our findings of decreased motivation for sucrose reward. However, it is unclear whether our change in corticosterone is an effect of habituation to the daily stress exposure or an effect of chronic stress on HPA-axis reactivity. It is important to note that, in the context of animal models of depression such as chronic social stress, corticosterone levels in females do not consistently predict behavioral changes [94]. Due to the limited availability of female rodent studies and the limited number of studies reporting corticosterone data, it is difficult to resolve the reasons behind the inconsistencies in rodent studies. It is unclear whether our results are an effect of this unique stressor and the ratio of witnessed versus experienced social defeat or the other factors discussed. Ultimately, this further underscores the importance of investigating effects of social stress on both physiological and behavioral responses in males and females.

## 5. Conclusion

The mechanisms by which stress alters motivated behaviors and thus promotes and exacerbates disorders of motivation are currently unresolved. We lack thorough characterization of social stress-induced changes in motivation with respect to time, sex, and stressor type. The present study sought to present a method by which we can begin to address the limited time points in which we monitor motivated behaviors. In doing so we also further supported the utility of a newer social defeat model that was inclusive of females. The consideration of sex as a biological factor revealed interesting divergent stress responses in males and females. These results demonstrate an exciting start to identifying the environmental, physiological, and neurobiological underpinnings of stress-induced disorders of motivation that are necessary for improved and inclusive treatment in humans.

## Acknowledgements

We thank Dr. Alexxai Kravitz and the members of the Feeding Experimentation Device (FED3) forum for guidance and support as we developed our protocol for continuous monitoring of operant responding in mice.

## References

[1] Hollon, N. G., Burgeno, L. M., Phillips, P. E. Stress effects on the neural substrates of motivated behavior. Nat Neurosci. 2015,18:1405–12.

[2] Dieterich, A., Stech, K., Srivastava, P., Lee, J., Sharif, A., Samuels, B. A. Chronic corticosterone shifts effort-related choice behavior in male mice. Psychopharmacology. 2020,237:2103–10.

[3] Stanton, C. H., Holmes, A. J., Chang, S. W. C., Joormann, J. From Stress to Anhedonia: Molecular Processes through Functional Circuits. Trends in neurosciences. 2019,42:23–42.

[4] Der-Avakian, A., Mazei-Robison, M. S., Kesby, J. P., Nestler, E. J., Markou, A. Enduring Deficits in Brain Reward Function after Chronic Social Defeat in Rats: Susceptibility, Resilience, and Antidepressant Response. Biological psychiatry. 2014,76:542–9.

[5] Dias-Ferreira, E., Sousa, J. C., Melo, I., Morgado, P., Mesquita, A. R., Cerqueira, J. J., et al. Chronic Stress Causes Frontostriatal Reorganization and Affects Decision-Making. Science. 2009,325:621–5.

[6] Pecoraro, N., Reyes, F., Gomez, F., Bhargava, A., Dallman, M. F. Chronic Stress Promotes Palatable Feeding, which Reduces Signs of Stress: Feedforward and Feedback Effects of Chronic Stress. Endocrinology. 2004,145:3754–62.

[7] Miczek, K. A., Nikulina, E. M., Shimamoto, A., Covinton, H. E. Escalated or Suppressed Cocaine Reward, Tegmental BDNF, and Accumbal Dopamine Caused by Episodic versus Continuous Social Stress in Rats \textbar Journal of Neuroscience. 2011.

[8] Sinha, R. Chronic Stress, Drug Use, and Vulnerability to Addiction. Ann N Y Acad Sci. 2008,1141:105–30.

[9] Naish, K. R., Laliberte, M., MacKillop, J., Balodis, I. M. Systematic review of the effects of acute stress in binge eating disorder. Eur J Neurosci. 2019,50:2415–29.

[10] Goldschmidt, A. B., Wonderlich, S. A., Crosby, R. D., Engel, S. G., Lavender, J. M., Peterson, C. B., et al. Ecological momentary assessment of stressful events and negative affect in bulimia nervosa. Journal of consulting and clinical psychology. 2014,82:30–9.

[11] Goddard, A. W. The Neurobiology of Panic: A Chronic Stress Disorder. Chronic Stress. 2017,1:2470547017736038.

[12] Conway, C. C., Rutter, L. A., Brown, T. A. Chronic Environmental Stress and the Temporal Course of Depression and Panic Disorder: A Trait-State-Occasion Modeling Approach. Journal of abnormal psychology. 2016,125:53–63.

[13] Atrooz, F., Alkadhi, K. A., Salim, S. Understanding stress: Insights from rodent models. Current Research in Neurobiology. 2021,2:100013.

[14] Canonica, T., Zalachoras, I. Motivational disturbances in rodent models of neuropsychiatric disorders. Frontiers in Behavioral Neuroscience. 2022,16.

[15] Buynitsky, T., Mostofsky, D. I. Restraint stress in biobehavioral research: Recent developments. Neurosci Biobehav Rev. 2009,33:1089–98.

[16] Eck, S. R., Bangasser, D. A. The effects of early life stress on motivated behaviors: A role for gonadal hormones. Neuroscience & Biobehavioral Reviews. 2020,119:86–100.

[17] Coccurello, R., D’Amato, F. R., Moles, A. Chronic social stress, hedonism and vulnerability to obesity: Lessons from Rodents. Neuroscience & Biobehavioral Reviews. 2009,33:537–50.

[18] Riga, D., Theijs, J. T., De Vries, T. J., Smit, A. B., Spijker, S. Social defeat-induced anhedonia: effects on operant sucrose-seeking behavior. Front. Behav. Neurosci. 2015,9.

[19] Bartolomucci, A., Cabassi, A., Govoni, P., Ceresini, G., Cero, C., Berra, D., et al. Metabolic Consequences and Vulnerability to Diet-Induced Obesity in Male Mice under Chronic Social Stress. PLOS ONE. 2009,4:e4331.

[20] Patterson, Z. R., Khazall, R., MacKay, H., Anisman, H., Abizaid, A. Central Ghrelin Signaling Mediates the Metabolic Response of C57BL/6 Male Mice to Chronic Social Defeat Stress. Endocrinology. 2013,154:1080–91.

[21] Smith, A., Hyland, L., Al-Ansari, H., Watts, B., Silver, Z., Wang, L., et al. Metabolic, neuroendocrine and behavioral effects of social defeat in male and female mice using the chronic non-discriminatory social defeat stress model. Hormones and Behavior. 2023,155:105412.

[22] Finger, B. C., Dinan, T. G., Cryan, J. F. The temporal impact of chronic intermittent psychosocial stress on high-fat diet-induced alterations in body weight. Psychoneuroendocrinology. 2012,37:729–41.

[23] Kleen, J. K., Sitomer, M. T., Killeen, P. R., Conrad, C. D. Chronic stress impairs spatial memory and motivation for reward without disrupting motor ability and motivation to explore. Behavioral Neuroscience. 2006,120:842–51.

[24] Dallman, M. F., Pecoraro, N. C., la Fleur, S. E. Chronic stress and comfort foods: self-medication and abdominal obesity. Brain, Behavior, and Immunity. 2005,19:275–80.

[25] Ulrich-Lai, Y. M., Ostrander, M. M., Thomas, I. M., Packard, B. A., Furay, A. R., Dolgas, C. M., et al. Daily limited access to sweetened drink attenuates hypothalamic-pituitary-adrenocortical axis stress responses. Endocrinology. 2007,148:1823–34.

[26] Shimamoto, A., Holly, E. N., Boyson, C. O., DeBold, J. F., Miczek, K. A. Individual differences in anhedonic and accumbal dopamine responses to chronic social stress and their link to cocaine self-administration in female rats \textbar Psychopharmacology. 2015.

[27] Barthas, F., Hu, M. Y., Siniscalchi, M. J., Ali, F., Mineur, Y. S., Picciotto, M. R., et al. Cumulative Effects of Social Stress on Reward-Guided Actions and Prefrontal Cortical Activity. Biological Psychiatry. 2020,88:541–53.

[28] Iio, W., Tokutake, Y., Matsukawa, N., Tsukahara, T., Chohnan, S., Toyoda, A. Anorexic behavior and elevation of hypothalamic malonyl-CoA in socially defeated rats. Biochemical and Biophysical Research Communications. 2012,421:301–4.

[29] Patterson, Z. R., Abizaid, A. B. Stress induced obesity: lessons from rodent models of stress. Front. Neurosci. 2013,7.

[30] Dieterich, A., Liu, T., Samuels, B. A. Chronic non-discriminatory social defeat stress reduces effort-related motivated behaviors in male and female mice. Transl Psychiatry. 2021,11:1–12.

[31] Bergamini, G., Cathomas, F., Auer, S., Sigrist, H., Seifritz, E., Patterson, M., et al. Mouse psychosocial stress reduces motivation and cognitive function in operant reward tests: A model for reward pathology with effects of agomelatine. European Neuropsychopharmacology. 2016,26:1448–64.

[32] Kúkel’ová, D., Bergamini, G., Sigrist, H., Seifritz, E., Hengerer, B., Pryce, C. R. Chronic Social Stress Leads to Reduced Gustatory Reward Salience and Effort Valuation in Mice. Front. Behav. Neurosci. 2018,12.

[33] Madur, L., Ineichen, C., Bergamini, G., Greter, A., Poggi, G., Cuomo-Haymour, N., et al. Stress deficits in reward behaviour are associated with and replicated by dysregulated amygdala-nucleus accumbens pathway function in mice. Commun Biol. 2023,6:1–20.

[34] Yoshida, K., Drew, M. R., Kono, A., Mimura, M., Takata, N., Tanaka, K. F. Chronic social defeat stress impairs goal-directed behavior through dysregulation of ventral hippocampal activity in male mice. Neuropsychopharmacol. 2021,46:1606–16.

[35] Bergamini, G., Mechtersheimer, J., Azzinnari, D., Sigrist, H., Buerge, M., Dallmann, R., et al. Chronic social stress induces peripheral and central immune activation, blunted mesolimbic dopamine function, and reduced reward-directed behaviour in mice. Neurobiol Stress. 2018,8:42–56.

[36] Lemon, C., Del Arco, A. Intermittent social stress produces different short- and long-term effects on effort-based reward-seeking behavior. Behavioural Brain Research. 2022,417:113613.

[37] Shaham, Y., Erb, S., Stewart, J. Stress-induced relapse to heroin and cocaine seeking in rats: a review. Brain Res Brain Res Rev. 2000,33:13–33.

[38] Newman, E. L., Leonard, M. Z., Arena, D. T., de Almeida, R. M. M., Miczek, K. A. Social defeat stress and escalation of cocaine and alcohol consumption: Focus on CRF. Neurobiol Stress. 2018,9:151–65.

[39] Blanchard, R. J., McKittrick, C. R., Blanchard, D. C. Animal models of social stress: effects on behavior and brain neurochemical systems. Physiology & Behavior. 2001,73:261–71.

[40] Miczek, K. A., Yap, J. J., Covington, H. E. Social stress, therapeutics and drug abuse: Preclinical models of escalated and depressed intake. Pharmacology & Therapeutics. 2008,120:102–28.

[41] Golden, S. A., Covington, H. E., Berton, O., Russo, S. J. A standardized protocol for repeated social defeat stress in mice. Nature protocols. 2011,6:1183–91.

[42] Newman, E. L., Covington, H. E., Suh, J., Bicakci, M. B., Ressler, K. J., DeBold, J. F., et al. Fighting females: Neural and behavioral consequences of social defeat stress in female mice. Biological psychiatry. 2019,86:657–68.

[43] Bartolomucci, A., Palanza, P., Gaspani, L., Limiroli, E., Panerai, A. E., Ceresini, G., et al. Social status in mice: behavioral, endocrine and immune changes are context dependent. Physiology & Behavior. 2001,73:401–10.

[44] Yohn, C. N., Dieterich, A., Bazer, A. S., Maita, I., Giedraitis, M., Samuels, B. A. Chronic non-discriminatory social defeat is an effective chronic stress paradigm for both male and female mice. Neuropsychopharmacology. 2019,44:2220–9.

[45] Azzinnari, D., Sigrist, H., Staehli, S., Palme, R., Hildebrandt, T., Leparc, G., et al. Mouse social stress induces increased fear conditioning, helplessness and fatigue to physical challenge together with markers of altered immune and dopamine function. Neuropharmacology. 2014,85:328–41.

[46] Kudryavtseva, N. N., Bakshtanovskaya, I. V., Koryakina, L. A. Social model of depression in mice of C57BL/6J strain. Pharmacology Biochemistry and Behavior. 1991,38:315–20.

[47] Furman, O., Tsoory, M., Chen, A. Differential chronic social stress models in male and female mice. Eur J Neurosci. 2022,55:2777–93.

[48] Lin, E. J., Sun, M., Choi, E. Y., Magee, D., Stets, C. W., During, M. J. Social overcrowding as a chronic stress model that increases adiposity in mice. Psychoneuroendocrinology. 2015,51:318–30.

[49] Sapolsky, R. M. Stress and the brain: individual variability and the inverted-U. Nat Neurosci. 2015,18:1344–6.

[50] Kuske, J. X., Trainor, B. C. Mean Girls: Social Stress Models for Female Rodents. Curr Top Behav Neurosci. 2022,54:95–124.

[51] Takahashi, A., Chung, J.-R., Zhang, S., Zhang, H., Grossman, Y., Aleyasin, H., et al. Establishment of a repeated social defeat stress model in female mice. Sci Rep. 2017,7:12838.

[52] Harris, A. Z., Atsak, P., Bretton, Z. H., Holt, E. S., Alam, R., Morton, M. P., et al. A Novel Method for Chronic Social Defeat Stress in Female Mice. Neuropsychopharmacol. 2018,43:1276–83.

[53] Bourke, C. H., Neigh, G. N. Exposure to repeated maternal aggression induces depressive-like behavior and increases startle in adult female rats. Behav Brain Res. 2012,227:270–5.

[54] Sial, O. K., Warren, B. L., Alcantara, L. F., Parise, E. M., Bolanos-Guzman, C. A. Vicarious social defeat stress: Bridging the gap between physical and emotional stress. J Neurosci Methods. 2016,258:94–103.

[55] Ródenas-González, F., Arenas, M. C., Blanco-Gandía, M. C., Manzanedo, C., Rodríguez-Arias, M. Vicarious Social Defeat Increases Conditioned Rewarding Effects of Cocaine and Ethanol Intake in Female Mice. Biomedicines. 2023,11:502.

[56] Boyson, C. O., Holly, E. N., Shimamoto, A., Albrechet-Souza, L., Weiner, L. A., DeBold, J. F., et al. Social Stress and CRF–Dopamine Interactions in the VTA: Role in Long-Term Escalation of Cocaine Self-Administration. J. Neurosci. 2014,34:6659–67.

[57] Engeln, M., Fox, M. E., Lobo, M. K. Housing conditions during self-administration determine motivation for cocaine in mice following chronic social defeat stress. Psychopharmacology. 2021,238:41–54.

[58] Yau, Y. H. C., Potenza, M. N. Stress and Eating Behaviors. Minerva endocrinologica. 2013,38:255–67.

[59] Ali, M. A., Kravitz, A. V. Challenges in Quantifying Food Intake in Rodents. Brain research. 2018,1693:188–91.

[60] Nguyen, K. P., O’Neal, T. J., Bolonduro, O. A., White, E., Kravitz, A. V. Feeding Experimentation Device (FED): A flexible open-source device for measuring feeding behavior. J Neurosci Methods. 2016,267:108–14.

[61] Mourra, D., Gnazzo, F., Cobos, S., Beeler, J. A. Striatal Dopamine D2 Receptors Regulate Cost Sensitivity and Behavioral Thrift. Neuroscience. 2020,425:134–45.

[62] Matikainen-Ankney, B. A., Earnest, T., Ali, M., Casey, E., Wang, J. G., Sutton, A. K., et al. An open-source device for measuring food intake and operant behavior in rodent home-cages. Elife. 2021,10.

[63] Grimm, J. W., North, K., Hopkins, M., Jiganti, K., McCoy, A., Šulc, J., et al. Sex differences in sucrose reinforcement in Long-Evans rats. Biology of Sex Differences. 2022,13:3.

[64] Radke, A. K., Sneddon, E. A., Monroe, S. C. Studying Sex Differences in Rodent Models of Addictive Behavior. Current protocols. 2021,1:e119.

[65] Ebner, K., Singewald, N. Individual differences in stress susceptibility and stress inhibitory mechanisms. Current Opinion in Behavioral Sciences. 2017,14:54–64.

[66] Wells, A. M., Ridener, E., Bourbonais, C. A., Kim, W., Pantazopoulos, H., Carroll, F. I., et al. Effects of Chronic Social Defeat Stress on Sleep and Circadian Rhythms Are Mitigated by Kappa-Opioid Receptor Antagonism. J. Neurosci. 2017,37:7656–68.

[67] Pawlyk, A. C., Morrison, A. R., Ross, R. J., Brennan, F. X. Stress-Induced Changes in Sleep in Rodents: Models and Mechanisms. Neurosci Biobehav Rev. 2008,32:99–117.

[68] Ding, G., Gong, Y., Eckel-Mahan, K. L., Sun, Z. Central Circadian Clock Regulates Energy Metabolism. Advances in experimental medicine and biology. 2018,1090:79–103.

[69] Acosta, J., Bussi, I. L., Esquivel, M., Höcht, C., Golombek, D. A., Agostino, P. V. Circadian modulation of motivation in mice. Behavioural Brain Research. 2020,382:112471.

[70] Spierling, S. R., Mattock, M., Zorrilla, E. P. Modeling hypohedonia following repeated social defeat: Individual vulnerability and dopaminergic involvement. Physiology & Behavior. 2017,177:99–106.

[71] Egan, A. E., Seemiller, L. R., Packard, A. E. B., Solomon, M. B., Ulrich-Lai, Y. M. Palatable food reduces anxiety-like behaviors and HPA axis responses to stress in female rats in an estrous-cycle specific manner. Hormones and Behavior. 2019,115:104557.

[72] Ulrich-Lai, Y. M., Christiansen, A. M., Ostrander, M. M., Jones, A. A., Jones, K. R., Choi, D. C., et al. Pleasurable behaviors reduce stress via brain reward pathways. Proceedings of the National Academy of Sciences. 2010,107:20529–34.

[73] Bangasser, D. A., Valentino, R. J. Sex differences in stress-related psychiatric disorders: neurobiological perspectives. Front Neuroendocrinol. 2014,35:303–19.

[74] Bezerra, H. S., Alves, R. M., Nunes, A. D. D., Barbosa, I. R. Prevalence and Associated Factors of Common Mental Disorders in Women: A Systematic Review. Public Health Rev. 2021,42:1604234.

[75] Kessler, R. C., McGonagle, K. A., Swartz, M., Blazer, D. G., Nelson, C. B. Sex and depression in the National Comorbidity Survey. I: Lifetime prevalence, chronicity and recurrence. Journal of Affective Disorders. 1993,29:85–96.

[76] Ramikie, T. S., Ressler, K. J. Mechanisms of Sex Differences in Fear and Posttraumatic Stress Disorder. Biol Psychiatry. 2018,83:876–85.

[77] Maeng, L. Y., Milad, M. R. Sex differences in anxiety disorders: Interactions between fear, stress, and gonadal hormones. Horm Behav. 2015,76:106–17.

[78] Altemus, M., Sarvaiya, N., Neill Epperson, C. Sex differences in anxiety and depression clinical perspectives. Front Neuroendocrinol. 2014,35:320–30.

[79] Khan, S. S., Secades-Villa, R., Okuda, M., Wang, S., Pérez-Fuentes, G., Kerridge, B. T., et al. Gender differences in cannabis use disorders: results from the National Epidemiologic Survey of Alcohol and Related Conditions. Drug and Alcohol Dependence. 2013,130:101–8.

[80] Vasilenko, S. A., Evans-Polce, R. J., Lanza, S. T. Age trends in rates of substance use disorders across ages 18-90: Differences by gender and race/ethnicity. Drug Alcohol Depend. 2017,180:260–4.

[81] Fonseca, F., Robles-Martínez, M., Tirado-Muñoz, J., Alías-Ferri, M., Mestre-Pintó, J.-I., Coratu, A. M., et al. A Gender Perspective of Addictive Disorders. Current Addiction Reports. 2021,8:89–99.

[82] Becker, J. B., Koob, G. F. Sex Differences in Animal Models: Focus on Addiction. Pharmacol Rev. 2016,68:242–63.

[83] McHugh, R. K., Votaw, V. R., Sugarman, D. E., Greenfield, S. F. Sex and gender differences in substance use disorders. Clinical Psychology Review. 2018,66:12–23.

[84] Bosch, O. J. Maternal aggression in rodents: brain oxytocin and vasopressin mediate pup defence. Philosophical Transactions of the Royal Society B: Biological Sciences. 2013,368:20130085.

[85] Iñiguez, S. D., Flores-Ramirez, F. J., Riggs, L. M., Alipio, J. B., Garcia-Carachure, I., Hernandez, M. A., et al. Vicarious Social Defeat Stress Induces Depression-Related Outcomes in Female Mice. Biological Psychiatry. 2018,83:9–17.

[86] Martínez-Caballero, M. Á., Calpe-López, C., García-Pardo, M. P., Arenas, M. C., de la Rubia Ortí, J. E., Bayona-Babiloni, R., et al. Behavioural traits related with resilience or vulnerability to the development of cocaine-induced conditioned place preference after exposure of female mice to vicarious social defeat. Progress in Neuro-Psychopharmacology and Biological Psychiatry. 2024,130:110912.

[87] Carnevali, L., Montano, N., Tobaldini, E., Thayer, J. F., Sgoifo, A. The contagion of social defeat stress: Insights from rodent studies. Neurosci Biobehav Rev. 2020,111:12–8.

[88] Warren, B. L., Mazei-Robison, M. S., Robison, A. J., Iñiguez, S. D. Can I Get a Witness? Using Vicarious Defeat Stress to Study Mood-Related Illnesses in Traditionally Understudied Populations. Biological Psychiatry. 2020,88:381–91.

[89] Nakatake, Y., Furuie, H., Yamada, M., Kuniishi, H., Ukezono, M., Yoshizawa, K., et al. The effects of emotional stress are not identical to those of physical stress in mouse model of social defeat stress. Neurosci Res. 2020,158:56–63.

[90] Macedo, G. C., Morita, G. M., Domingues, L. P., Favoretto, C. A., Suchecki, D., Quadros, I. M. H. Consequences of continuous social defeat stress on anxiety- and depressive-like behaviors and ethanol reward in mice. Hormones and Behavior. 2018,97:154–61.

[91] McQuaid, R. J., Audet, M.-C., Jacobson-Pick, S., Anisman, H. The differential impact of social defeat on mice living in isolation or groups in an enriched environment: plasma corticosterone and monoamine variations. International Journal of Neuropsychopharmacology. 2013,16:351–63.

[92] Heim, C., Ehlert, U., Hellhammer, D. H. The potential role of hypocortisolism in the pathophysiology of stress-related bodily disorders. Psychoneuroendocrinology. 2000,25:1–35.

[93] Marchette, R. C. N., Bicca, M. A., Santos, E., de Lima, T. C. M. Distinctive stress sensitivity and anxiety-like behavior in female mice: Strain differences matter. Neurobiol Stress. 2018,9:55–63.

[94] Kokras, N., Krokida, S., Varoudaki, T. Z., Dalla, C. Do corticosterone levels predict female depressive-like behavior in rodents? J Neurosci Res. 2021,99:324–31.

